# Impact of Mutations Affecting 4’-Fluorouridine Susceptibility on Fitness and Treatment Outcomes for Venezuelan Equine Encephalitis Virus

**DOI:** 10.1101/2025.09.11.675625

**Authors:** Jenny Wong, Tahirah Moore, Devin Lewis, Carolin M Lieber, Dariia Vyshenska, Elizabeth B. Sobolik, Sarah Zimmermann, George R Painter, Richard K Plemper, Alexander L Greninger, Robert M Cox

## Abstract

Venezuelan equine encephalitis virus (VEEV) is a prototypical encephalitic alphavirus. Members of the *Alphavirus* genus are found across the globe, transmitted by arthropod vectors, and cause significant disease burdens in humans and animals. There are currently no FDA-approved antivirals against any member of the *Alphavirus* genus. While a vaccine exists against chikungunya virus (CHIKV), a member of the arthitogenic alphaviruses, FDA-approved vaccines are not available for other members of this genus, particularly the encephalitic alphaviruses such as VEEV, Eastern equine encephalitis virus (EEEV) and Western equine encephalitis virus (WEEV). 4’-Fluorouridine (4’-FlU, EIDD-2749) was recently identified as a broad-spectrum antiviral against multiple RNA viruses, including alphaviruses. 4’-FlU can potently inhibit VEEV-TC83 replication, with submicromolar potency in cell culture. However, the emergence of antiviral resistance represents a hurdle for antiviral drug development and the implementation of effective treatment strategies. Here, we have identified novel mutations in the VEEV nsP4 RNA-dependent RNA polymerase that reduce susceptibility to 4’-FlU, including P187A, Q191L, L289F, and T296I. We rebuilt each mutation in recombinant VEEV-TC83 and characterized the effects of these mutations on fitness and pathogenicity. In addition, we assessed the impact of mutations reducing sensitivity to 4’-FlU in a mouse model. Although mutations against 4’-FlU arise quickly *in vitro*, treatment can still alleviate severe disease and lethal encephalitis. Together, these data highlight the promising therapeutic potential of 4’-FlU for the treatment of alphavirus encephalitis.

**Importance:** VEEV is one of several mosquito-spread viruses that can cause serious brain infections in people and animals. Unlike CHIKV, against which an approved vaccine exists, there are no countermeasures to prevent or treat VEEV infections or block its close relatives, EEEV and WEEV. 4’-FlU inhibits many different RNA viruses, including VEEV, and is currently being developed to treat multiple viral infections. A major problem with small-molecule antivirals is the appearance of virus populations that are less susceptible to treatment. In this study, we identified mutations in VEEV’s RNA-dependent RNA polymerase (nsP4) that confer reduced susceptibility to 4’-FlU. We then engineered these mutations into full-length infectious clones and assessed if viruses encoding for these mutations were still pathogenic. In addition, we treated infected mice with 4’-FlU and measured how well the compound inhibited virus replication and prevented severe disease, even when mice were infected with viruses harboring mutations that reduced susceptibility to 4’-FlU. Although VEEV can develop moderate resistance to 4’-FlU *in vitro*, administration of 4’-FlU still reduced severe disease and prevented lethality in the animals infected with viruses that possess mutations that decrease susceptibility to 4’-FlU. These results suggest that 4’-FlU has strong potential as a future treatment for alphavirus infections like VEEV encephalitis.

## Introduction

Alphaviruses are mosquito-borne, positive-strand RNA viruses that circulate on every inhabited continent. Alphaviruses can be partitioned into arthritogenic and encephalitic groups. Whereas arthitogenic members such as CHIKV and Mayaro virus predominantly cause self-limiting febrile arthritis (1–3), encephalitic alphaviruses, most notably VEEV, EEEV, and WEEV, invade the central nervous system and can be fatal (4, 5). VEEV, the most frequently encountered encephalitic species, is responsible for periodic outbreaks in the Americas, replicates to high levels in vertebrate and mosquito cells, and is classified as a potential biothreat agent because of its stability in aerosols and lack of licensed countermeasures (6).

The ∼11.5-12 kb alphavirus genome contains two open reading frames (7, 8). The 5′ ORF encodes the non-structural polyprotein (P1234), which is processed into nsP1-nsP4 to build the viral replication complex (9). Replication takes place inside membrane invaginations, where nsP1 forms a membrane-anchored dodecameric ring, nsP2 contributes helicase and protease activities, nsP3 recruits host factors, and nsP4 functions as the RNA-dependent RNA polymerase (RdRp) (10–14). Because the viral RdRp lacks a host cell-equivalent and is critical for virus replication, nsP4 represents an attractive target for direct-acting antivirals.

Multiple small-molecule antivirals have sought to exploit this vulnerability. Broad-spectrum nucleoside analogues such as ribavirin, β-D-N^4^-hydroxycytidine (NHC), favipiravir, and sofosbuvir inhibit CHIKV or VEEV replication by either reducing nucleotide pools or being incorporated into nascent RNA strands, causing premature termination or lethal mutagenesis (15–21). Mutations conferring reduced sensitivity to these agents consistently map to nsP4 (19, 22). For example, C483Y in CHIKV confers ribavirin and 5-fluorouracil tolerance (23), whereas P187S/A189V/I190T in VEEV were shown to reduce susceptibility to NHC (19).

4’-FlU is a next-generation uridine analogue that triggers immediate or delayed chain termination of viral RNA synthesis (24, 25). Originally developed against respiratory pathogens, including SARS-CoV-2, respiratory syncytial virus (25), and influenza virus (26), 4’-FlU has recently demonstrated nanomolar potency against CHIKV in cell culture and, following oral administration, has reduced viral load, tissue inflammation, and clinical disease in mice infected with CHIKV and Mayaro virus (27). Preliminary passaging studies with CHIKV identified single amino acid substitutions (Q192L or C483Y) in nsP4 that partially attenuate drug sensitivity without enhancing pathogenicity, and suggested that the compound possesses a relatively high genetic barrier against resistance (28). However, the 4’-FlU susceptibility profile of other alphavirus targets is undefined.

Given the public health importance of VEEV and the historical propensity of nucleoside analogues to select RdRp variants, defining the landscape of VEEV mutations associated with reduced susceptibility to 4’-FlU is a critical step in the pre-clinical development of the compound. Here, we characterize the capacity of VEEV to acquire mutations that can lead to reduced potency against 4’-FlU. Using serial passages under escalating 4’-FlU concentrations, we isolated VEEV variants with decreased susceptibility, mapped and rebuilt the responsible nsP4 mutations, and evaluated their effects on viral fitness, pathogenic potential, and treatment efficacy. These data provide a framework of sites within nsP4 that must be monitored during clinical deployment and inform therapeutic strategies for potential use of 4’-FlU against encephalitic alphaviruses.

## Results

### 4’-FlU exhibits broad-spectrum potency

The indication spectrum of 4’-FlU was first assessed in tissue culture, measuring effects on virus growth in a range of reporter virus assays. For reference, we repeated dose-response assays using several viruses previously shown to be inhibited by 4’-FlU (24, 25, 27). All positive-sense and negative-sense RNA viruses tested were inhibited by 4’-FlU. These assays showed that 4’-FlU can inhibit a range of positive-and negative-strand RNA viruses, including VEEV, CHIKV, Sindbis virus (SINV), dengue virus 2 (DENV2), SARS-CoV-2, human parainfluenza virus 3, Sendai virus (SeV), measles virus, Cedar virus, and vesicular stomatitis virus, with 50%-effective concentration (EC_50_) values ranging from 0.12-7.7 µM (**Figure 1**) (24, 25, 27). In our preliminary dose response testing, 4’-FlU was capable of inhibiting VEEV-TC83 with sub-micromolar potency (EC50 = 0.22 µM). Toxicity on VeroE6 cells showed minimal cytotoxicity, with a 50% cytotoxic concentration (CC_50_) of >100 µM.

**Figure 1.**
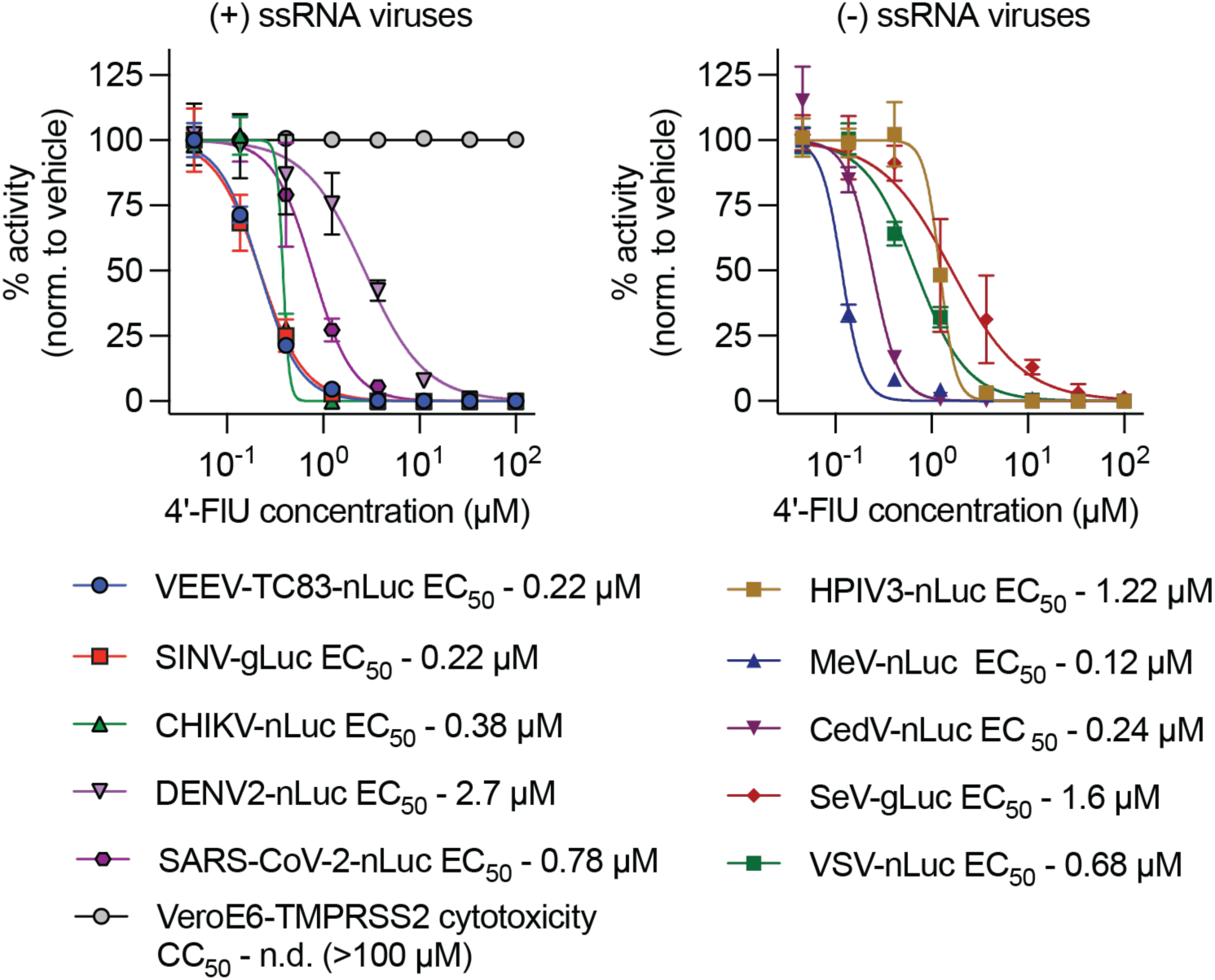
Indication spectrum of 4’-FlU. 4’-FlU has a broad activity spectrum covering both positive-(+) and negative-sense (-) single stranded RNA (ssRNA) viruses. 50% effective concentrations (EC_50_) are shown along with the 50% cytotoxic concentration (CC_50_) on VeroE6-TMPRSS2 cells. Assay results agree with previous reports of potency against various positive and negative sense RNA viruses (24, 25, 27). Symbols represent means of biological repeats (n≥3), lines intersect data means, and error bars represent standard deviation. EC_50_ concentrations were calculated through 4-parameter variable slope regression modeling.

### *In vitro* adaptation of VEEV-TC83 against 4’-FlU

To assess barriers of resistance and identify mutations that confer reduced susceptibility to 4’-FlU, VEEV-TC83 expressing GFP under a second subgenomic promoter (**Figure 2**) was passaged in the presence of increasing concentrations of 4’-FlU in BHK-21 cells (29). For each passage, cells were infected with a multiplicity of infection (MOI) of 0.5-1. To identify mutations that confer reduced susceptibility to 4’-FlU, 14 independent lineages of VEEV-eGFP were serially passaged in the presence of increasing concentrations of 4’-FlU (0.5-10 µM). As a control, 6 lineages of VEEV-eGFP were passaged in the presence of vehicle (DMSO volume equivalents). For each passage, virus was harvested 18-24 hours after infection, or at maximum cytopathic effect (CPE). Infections were monitored by visualizing GFP expression. After 5 passages, virus was harvested, and RNA was extracted for whole genome sequencing. VEEV-eGFP passaged in the presence of DMSO vehicle possessed no mutations with allele frequency >10%. In 4’-FlU-experienced lineages, multiple nsP4 mutations with allele frequency >10% were identified, including substitutions at the previously identified Q191L position (CHIKV Q192L) (28) and surprisingly, P187A, the location of a mutation associated with reduced susceptibility to β-D-N^4^-hydroxycytidine (NHC), the bioactive metabolite of molnupiravir (19), in VEEV-TC83. A panel of novel mutations in nsP4 furthermore emerged, including A146V, L289F, and T296I (**Figure 2**; summarized in **Table 1**). Mutated residues were highly conserved across the alphavirus genus (Figure 2). The only mutation not 100% conserved across species was L289F, which is present as a methionine in Old World alphaviruses. The spatial organization was assessed by mapping each nsP4 mutation identified in this study studies onto a homology model of the VEEV nsP4 polymerase (**Figure 2**). These mutations all clustered opposite of the GDD active site for phosphodiester bond formation within nsP4 core, likely allowing for altered RNA secondary structure induced by incorporation of 4’-FlU, similar to what has been described for influenza viruses (30).

**Figure 2.**
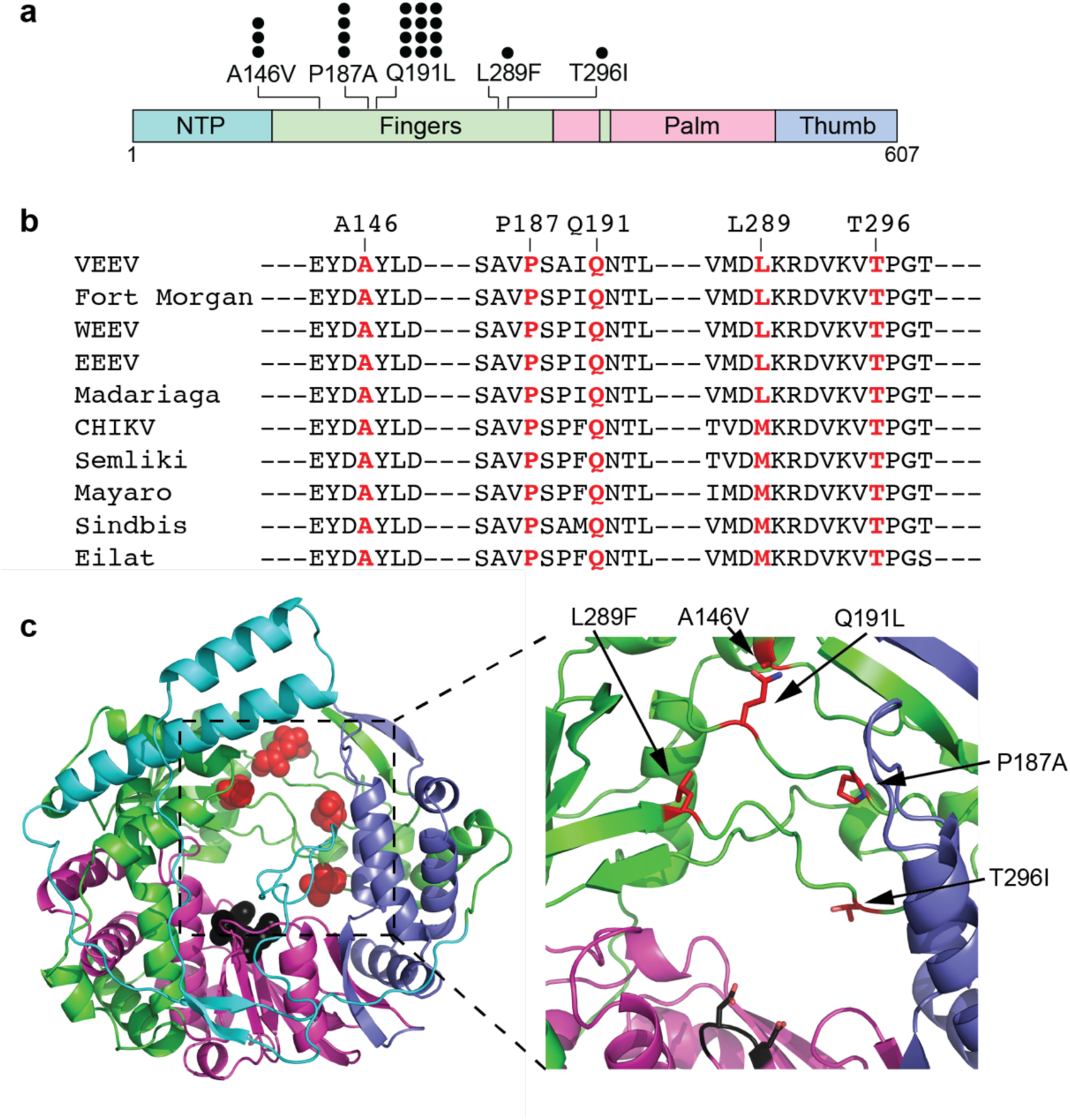
Susceptibility profiling studies. A) VEEV-TC83-eGFP was serially passaged (5 passages) in the presence of increasing concentrations of 4’-FlU. Mutations associated with increased concentrations of 4’-FlU were identified in nsP4. Each circle represents a lineage in which the mutation was identified (14 4’-FlU lineages; 6 DMSO lineages). No mutations above 10% frequency arose in DMSO passaged populations. B) Conservation of 4’-FlU associated mutations across the alphavirus genus. C) Spatial orientation of 4’-FlU associated mutations in a model of the VEEV nsP4 polymerase based on CHIKV nsP4 (PDB ID: 7Y38) (34). Red spheres or sticks denote residues identified in susceptibility profiling studies. Black spheres denote the GDD active site.

**Table 1.** Allele frequency of nsP4 mutations in VEEV-TC83 lineages.

|  |  | VEEV-TC83 lineage |  |  |  |  |  |  |  |  |  |  |  |  |  |
| --- | --- | --- | --- | --- | --- | --- | --- | --- | --- | --- | --- | --- | --- | --- | --- |
|  |  | 1 | 2 | 3 | 4 | 5 | 6 | 7 | 8 | 9 | 10 | 11 | 12 | 13 | 14 |
| *Mutation* | A146V | -- | -- | -- | -- | 48.1 | -- | -- | -- | -- | 73.9 | -- | 62.2 | -- | -- |
|  | L289F | -- | -- | -- | -- | -- | -- | -- | -- | 77.5 | -- | -- | -- | -- | -- |
|  | P187A | -- | 91.2 | 44.9 | -- | 13.2 | -- | -- | 16.4 | -- | -- | -- | -- | -- | -- |
|  | Q191L | 85 | -- | 37.1 | 87.3 | 26.8 | 99.5 | 95.7 | 80.2 | -- | 22.7 | 80.8 | 35.3 | 99.5 | 99 |
|  | T296I | -- | -- | -- | -- | -- | -- | -- | -- | -- | -- | -- | -- | -- | 20.8 |

nsP4 mutations with allele frequency >10% are shown.

### Effects of mutations on 4’-FlU susceptibility *in vitro*

To measure the effects of nsP4 substitutions on susceptibility to 4’-FlU, each mutation was rebuilt individually into a full-length VEEV-NLuc-PEST reporter virus, harboring a nano-luciferase reporter with C-terminal PEST sequence. After authentication, recombinant viruses were tested in dose-response assays for changes in sensitivity to 4’-FlU (**Figure 3**). To gauge the effects of each mutation on 4’-FlU susceptibilty, the parent VEEV-TC83 nLuc was tested in parallel to VEEV-TC83 nLuc constructs encoding for each mutation identified in susceptibility profling studies (**Figure 3**). Similar to previous dose response data (**Figure 1**), 4’-FlU potently inhibited VEEV-TC83 nLuc, with an EC50 of 0.19 µM (**Figure 3**). Of all constructs tested, only A146V showed no changes in susceptibility to the 4’-FlU. Interestingly, A146V is located in close proximity to Q191L and appeared only in lineages that also possessing the Q191L mutation, suggesting a possible compensatory role for A146V. All other mutations led to moderate decreases in 4’-FlU potency (∼3 to 6-fold increases in EC_50_, Table 2). No single mutation provided complete tolerance to 4’-FlU. Combination of the two most prevalent mutations, P187A and Q191L, provided the largest decrease in potency, with EC_50_ and EC_90_ values of 1.6 and 5.0 µM, respectively. To assess whether there is cross resistance with NHC, the clinically most advanced nucleoside analog with confirmed efficacy against alphaviruses, all mutations were tested against this inhibitor also. All constructs displayed similar susceptibility to NHC compared to parent VEEV-NlucPEST-TC83 (EC_50_ = 0.67 µM), with EC_50_ values ranging from 0.36-0.79 µM (**Figure 4**).

**Figure 3.**
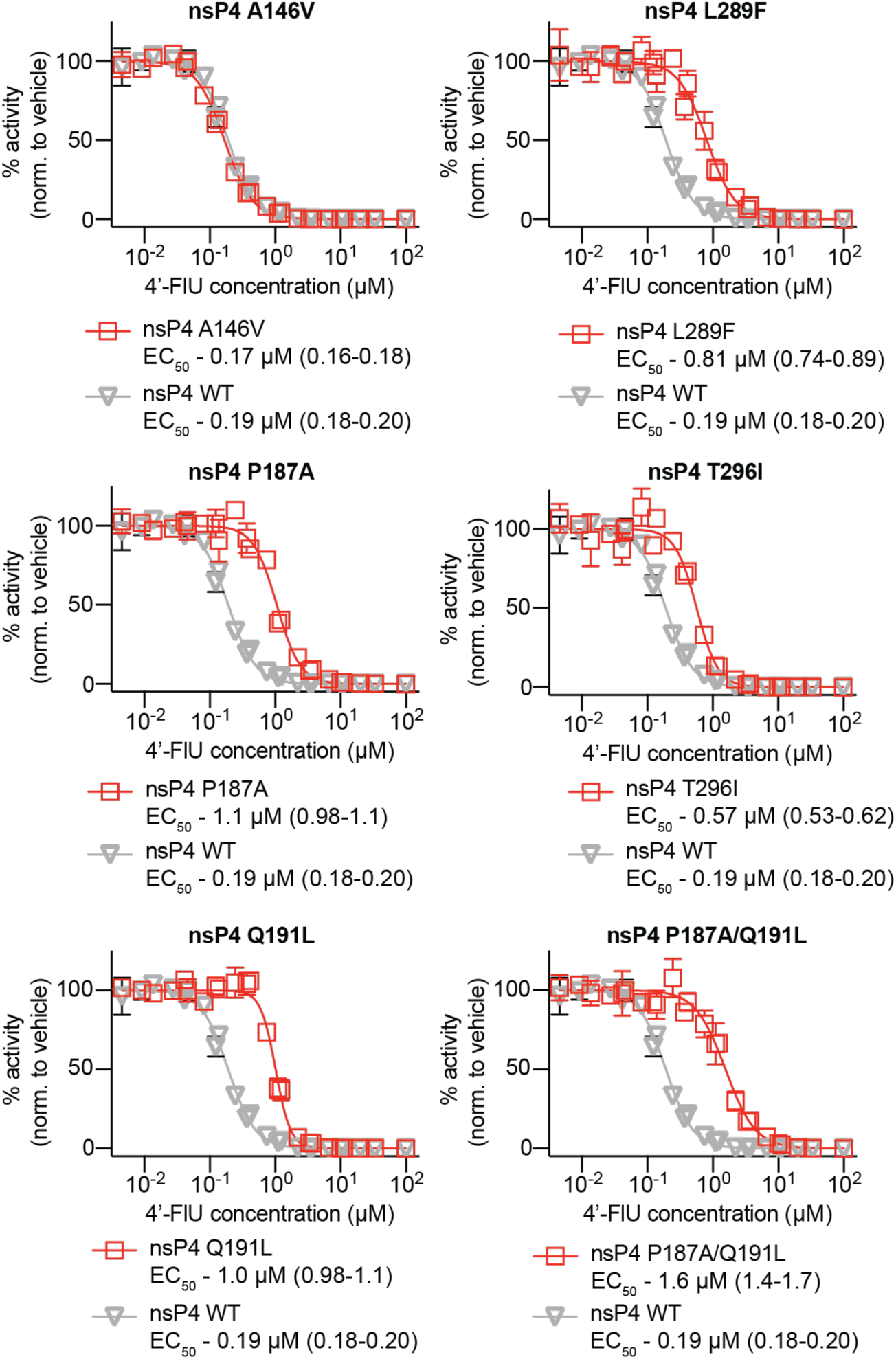
Effects of mutations on 4’-FlU susceptibility. Individual 4’-FlU mutations were rebuilt into full-length VEEV nLuc reporter viruses. Changes in potency were determined by dose-response assay. For each mutation tested, the parent (VEEV-TC83-nLuc) inhibition profile is shown (gray). EC_50_ values for each construct are shown along with confidence intervals in parentheses. Symbols represent means of biological repeats (n=3), lines intersect data means, and error bars represent standard deviation. EC_50_ concentrations were calculated through 4-parameter variable slope regression modeling.

**Figure 4.**
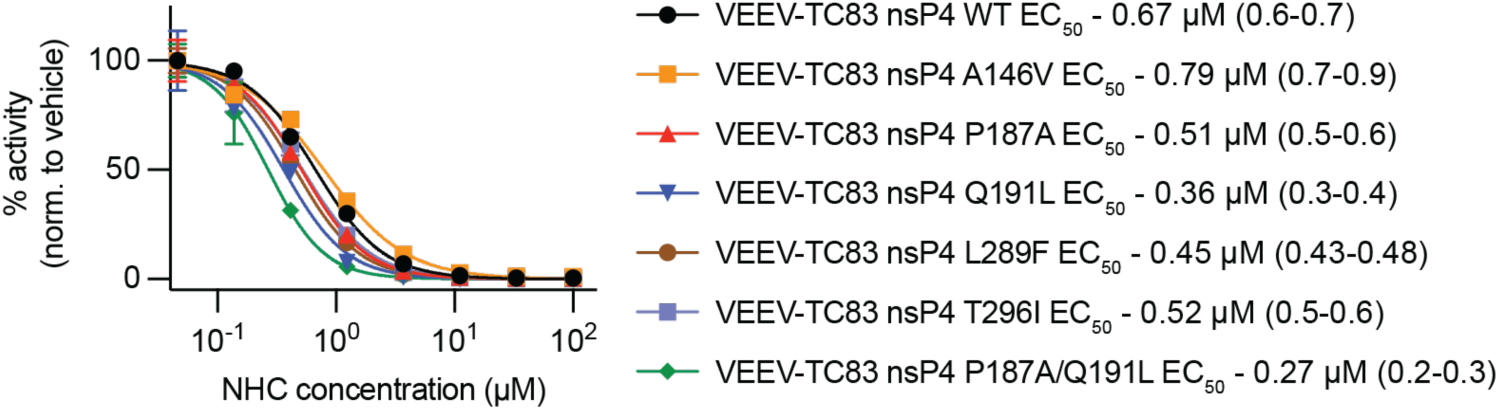
Effects of mutations on NHC susceptibility. For mutations associated with reduced susceptibility to 4’-FlU, potency was assessed against NHC by dose-response assay. No 4’-FlU associated mutation conferred reduced susceptibility to NHC. EC_50_ values for each construct are shown along with confidence intervals in parentheses. Symbols represent means of biological repeats (n=3), lines intersect data means, and error bars represent standard deviation. EC_50_ concentrations were calculated through 4-parameter variable slope regression modeling.

**Table 2.** EC_50_/EC_90_ fold change for each 4’-FlU associated mutation.

|  | EC<sub>50</sub> (μM) | Fold change | EC<sub>90</sub> (μM) | Fold change |
| --- | --- | --- | --- | --- |
| WT | 0.19 (0.18-0.2) | 1.00 | 0.55 (0.5-0.6) | 1.00 |
| A146V | 0.17 (0.16-0.18) | 0.88 | 0.50 (0.45-0.55) | 0.92 |
| P187A | 1.1 (0.99-1.1) | 5.51 | 2.6 (2.2-3.1) | 4.80 |
| Q191L | 1.0 (0.98-1.1) | 5.30 | 1.8 (1.7-2.0) | 3.33 |
| L289F | 0.81 (0.74-0.89) | 4.23 | 2.4 (2.0-3.0) | 4.46 |
| T296I | 0.58 (0.53-0.62) | 2.98 | 1.3 (1.1-1.5) | 2.39 |
| P187A/Q191L | 1.6 (1.4-1.7) | 8.13 | 5.0 (4.1-6.1) | 9.12 |

To determine if resistance mechanisms are shared across encephalitic alphavirus species, select mutations were engineered into an EEEV replicon (**Figure 5**). In the EEEV replicon, 4’-FlU was found to potently inhibit polymerase activity with an EC_50_ of 2.86 µM. Similar to VEEV, EEEV mutations P188A and Q192L led to moderate decreases in susceptibility to 4’-FlU, with EC_50_ values of 8.46 µM and 5.6 µM, respectively (**Figure 5**). These data, combined with previous reports for CHIKV, suggest a conserved mechanism for 4’-FlU tolerance across the *Alphavirus* genus.

**Figure 5.**
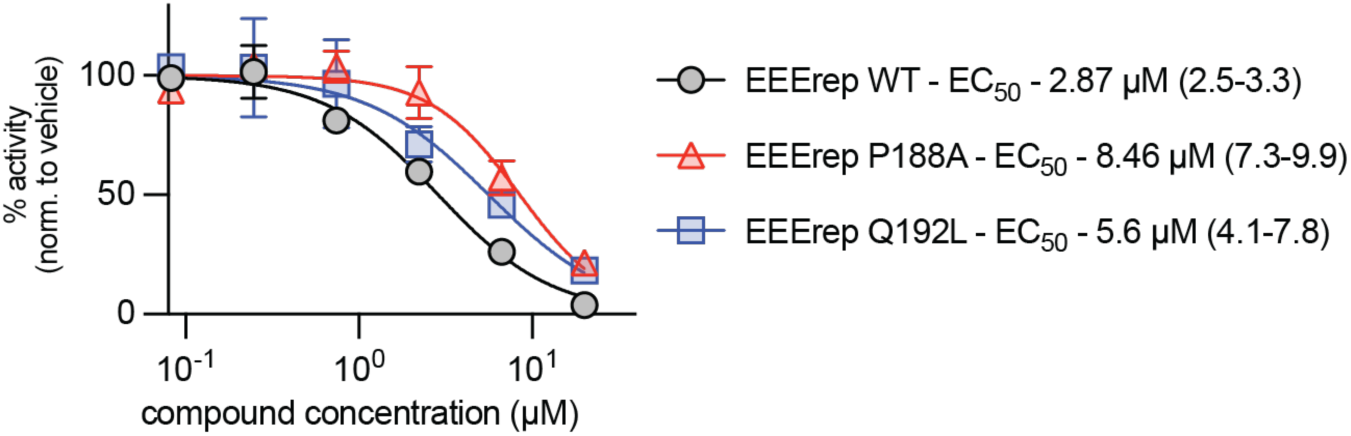
Effects of select mutations on 4’-FlU potency in an Eastern equine encephalitis virus replicon. The P188A and Q192L 4’-FlU mutations were rebuilt into a EEEV nLuc replicon. Changes in potency were determined by dose-response assay. The parent EEErep WT inhibition profile is shown (gray). EC_50_ values for each construct are shown along with confidence intervals in parentheses. Symbols represent means of biological repeats (n=3), lines intersect data means, and error bars represent standard deviation. EC_50_ concentrations were calculated through 4-parameter variable slope regression modeling.

### Effects of mutations on *in vitro* fitness

To assess the impact of mutations on VEEV-TC83 growth kinetics and fitness, growth curves were generated for mutated viruses in BHK-21 cells. Cells were infected with an MOI of 0.1. Supernatant was harvested at multiple time points after infection and subjected to TCID_50_ titration. Recombinant viruses encoding for P187A, A146V, L289F, or T296I displayed significantly lower titers at one or more time points preceding 24 hours after infection, suggesting a negative impact on growth kinetics (**Figure 6**). VEEV nsP4 encoding for a Q191L mutation did not display any significant reductions in growth. VEEV expressing the combination of P187A and Q191L mutations exhibited a significantly slower growth kinetics compared to parent VEEV-TC83-nLucPEST (**Figure 4**). All variants reached peak titers of 10^8^-10^9^ TCID_50_/ml between 18-36 hours post-infection.

**Figure 6.**
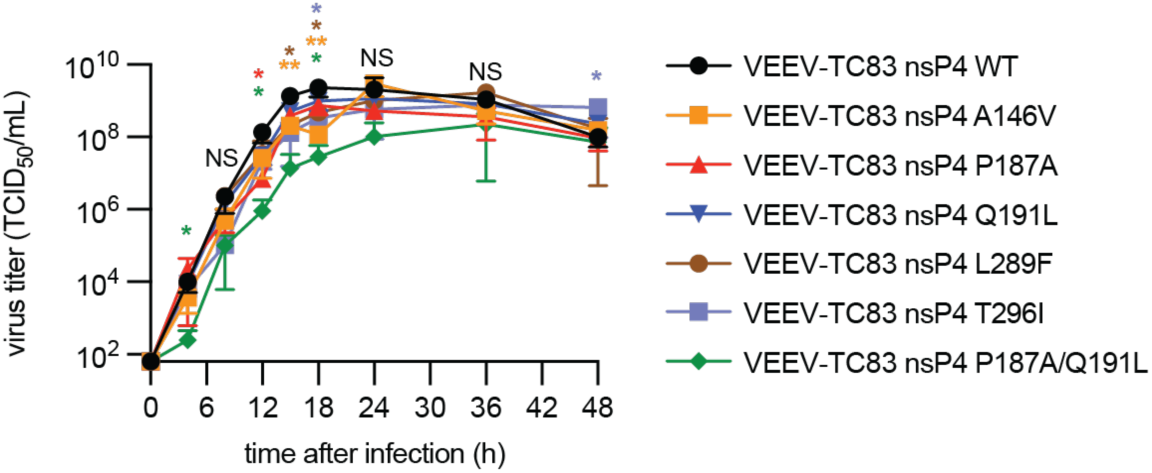
Growth curves of standard VEEV-TC83 and VEEV-TC83 with potential 4’-FlU associated mutations in BHK-21 cells. Cells were infected (MOI=0.1), and supernatant was harvested at multiple time points after infection. Virus titer was determined by TCID_50_ titration. Symbols represent biological repeats (n=3), lines indicate means, and error bars show SD. Statistical analysis through two-way ANOVA and Dunnett’s multiple comparisons post-hoc test. Significance is defined by the following P value ranges: P>0.05 (NS), P≤0.05 (*), P≤0.01 (**), P≤0.001 (***), P≤0.0001 (****).

### Effects of 4’-FlU associated mutations on *in vivo* pathogenesis and treatment

To determine the effects of mutations conferring reduced susceptibility to 4’-FlU on *in vivo* pathogenesis and assess the effect of treatment in the context of mutant virus populations, C3H/HeN mice were infected with 1 × 10^6^ TCID_50_ of VEEV-TC83 (n=6) or VEEV-TC83 encoding one of the following mutations: P187A (n=8), Q191L (n=8), L289F (n=8), and P187A/Q191L (n=8) (**Figure 7**). Infected mice were treated with vehicle or 4’-FlU (5 mg/kg; q.d.; starting 12 hours after infection). Since P187 has been associated with reduced susceptibility to NHC/molnupiravir, an additional group of mice infected with the VEEV-TC83 nsP4-P187A was treated with molnupiravir (150 mg/kg; b.i.d.) (31). Subgroups of mice (n=3) were euthanized 4 days after infection to assess viral loads in the brain. Bodyweight and clinical signs were monitored daily.

**Figure 7.**
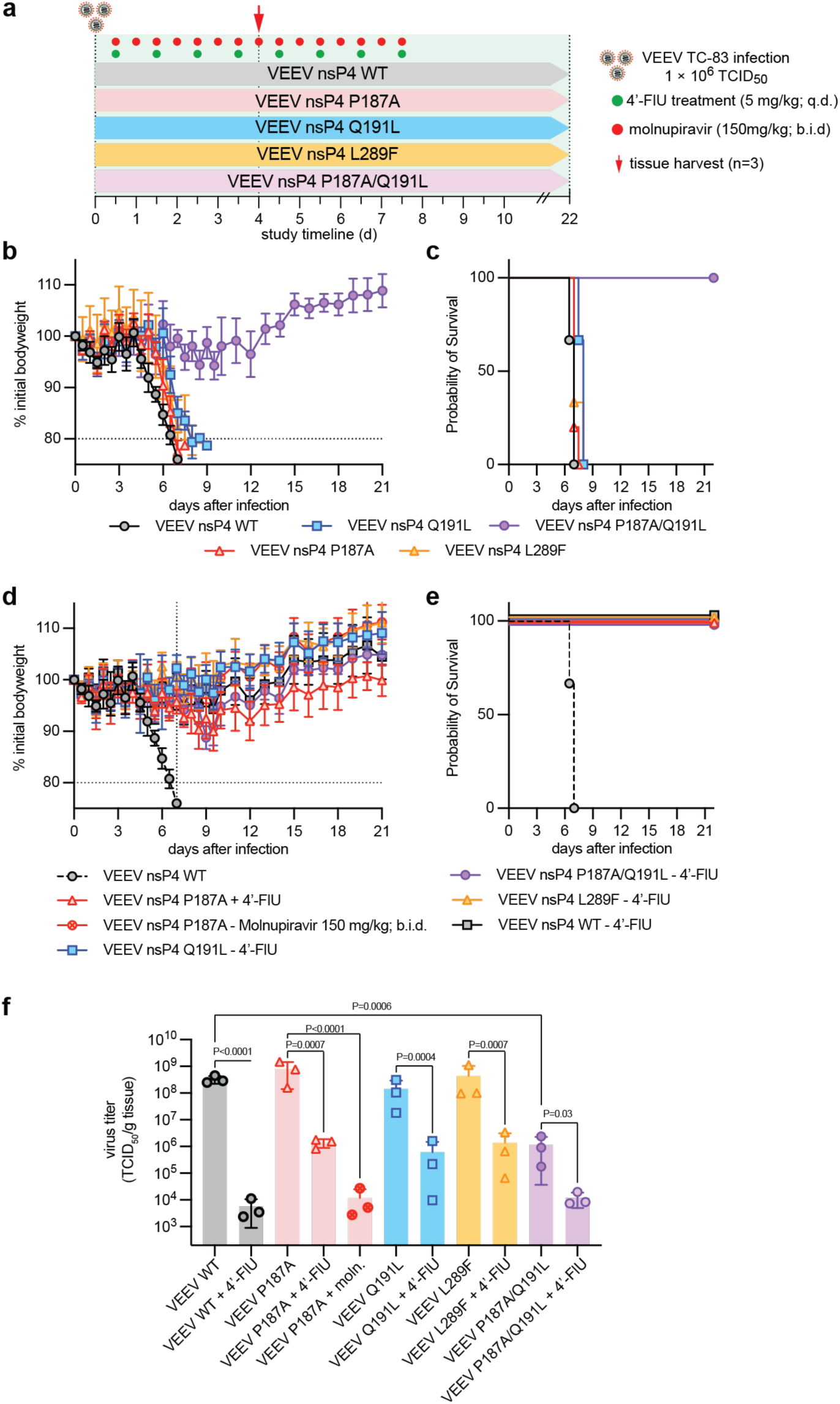
Effect of 4’-FlU associated mutations on *in vivo* efficacy, and correlation between 4’-FlU susceptibility and viral pathogenesis. A) Schematic of 4’-FlU efficacy and pathogenesis study. Animals were treated therapeutically with 4’-FlU at 5 mg/kg dose concentration, administered once daily (q.d.). B-C) C3H/HeN mice (n=8) were infected with 1 × 10^6^ TCID_50_ units of VEEV-TC83 or VEEV-TC83 encoding for individual 4’-FlU associated mutations. Clinical signs (body weight) were monitored daily (b). Survival (c) was calculated 21 days after infection. Survival analysis was performed using time-to-event log-rank (Mantel-Cox) test. D-E) Effect of therapeutic treatment with 4’-FlU on disease outcome. C3H/HeN mice (n=8) were infected with 1 × 10^6^ TCID_50_ units of VEEV-TC83 or VEEV-TC83 encoding for individual 4’-FlU associated mutations. Treatment was initiated 12 hours after infection with 4’-FlU (5 mg/kg dose; q.d.) and continued for 7 days. F) Four days after infection, brain tissue was harvested from subgroups of mice (n=3) to assess viral load. Symbols represent individual animals. Error bars represent standard deviation. Bars represent data means. Statistical analysis through one-way ANOVA with Tukey’s multiple comparisons post-hoc test. Significance is defined by the following P value ranges: P>0.05 (NS), P≤0.05 (*), P≤0.01 (**), P≤0.001 (***), P≤0.0001 (****).

Vehicle-treated mice infected with all VEEV-TC83 variants, with the exception of nsP4 P187/Q191L, reached a predefined endpoint (>20% body weight loss) by 9 days post-infection. Vehicle-treated mice infected with VEEV-TC83, VEEV-TC83 nsP4-P187A, and VEEV-TC83 nsP4-L289F had a median survival of 7 days, whereas mice infected with VEEV-TC83 harboring nsP4-Q191L experienced a median survival of 8 days. Infection with a all constructs, except the P187A/Q191L variant, led to the appearance of clinical signs approximately 5-6 days after infection, including loss of body weight, aggression, and restlessness. None of the mice infected with the P187A/Q191L variant reached endpoint (>20% loss in body weight) and clinical signs displayed were minimal. Treatment of infected mice with 4’-FlU prevented the appearance of any clinical signs, provided complete protection from lethal disease, and led to significant reductions in virus load in the brain of infected mice across all variants tested. In addition, mice infected with the P187A variant and treated with molnupiravir survived until study end and showed significant reductions in virus load by day 4. Treatment led to substantial decreases in bodyweight loss.

## Discussion

Here we demonstrate that 4’-FlU potently inhibits VEEV replication both *in vitro* and *in vivo*, consistent with previously reported activity against other alphaviruses, including CHIKV (27). While mutations against 4’-FlU have been described in influenza virus (30) and CHIKV, our study provides the first in-depth characterization of 4’-FlU tolerance in VEEV, an archetype encephalitic alphavirus. Using serial passaging of VEEV-TC83 in the presence of 4’-FlU, we identified five 4’-FlU-associated mutations in the viral nsP4 RdRp: the previously reported Q191L, and four novel mutations, A146V, P187A, L289F, and T296I.

Due to the high mutation rate compared to other RNA viruses, and lack of proofreading in alphavirus replication, nsP4 variants emerged rapidly after only five passages (32, 33). All substitutions associated with 4’-FlU passaging were located in the nsP4 polymerase and mapped to the fingers domain, opposite the catalytic GDD motif, in a homology model of VEEV nsP4 based on the CHIKV nsP4 structure (PDB ID: 7Y38) (**Figure 2**) (34). Notably, these residues are positioned within the central pore of the polymerase, distant from contacts with other nonstructural proteins, suggesting a potential role in RNA template recognition or positioning during synthesis. This spatial localization supports a mechanism in which the identified mutations may alter the ability of nsP4 to engage RNA following incorporation of 4’-FlU, similar to models proposed for 4’-FlU tolerance in influenza virus (30). However, in the absence of high-resolution RNA-bound structures of alphavirus nsP4, definitive mechanistic interpretations remain speculative.

Using reverse genetics, we engineered each mutation individually into the VEEV-TC83 backbone to assess its effect on replication fitness and susceptibility to 4’-FlU. Similar to CHIKV nsP4 variants, VEEV mutations conferred modest reductions in viral fitness and resulted in only moderate decreases in potency (3-to 5-fold increases in EC_50_), with none conferring complete tolerance to 4’-FlU. Interestingly, the A146V mutation had no impact on drug susceptibility and was only observed in lineages that also contained Q191L, suggesting a potential compensatory interaction between these adjacent residues (**Figure 2**). In contrast to CHIKV, VEEV did not emerge with the fidelity-enhancing C483Y mutation in nsP4, highlighting species-specific differences in the evolution of nucleoside analog resistance (28, 35).

Notably, the Q191L and P187A mutations appear to represent conserved determinants across alphaviruses. P187, in particular, has now been associated with reduced susceptibility to both 4’-FlU and NHC, two nucleoside analogs with distinct mechanisms of action, chain termination (4’-FlU) versus lethal mutagenesis (NHC), respectively. This dual tolerance profile underscores the functional relevance of this residue in nucleobase recognition. However, despite its association with NHC tolerance (P187S), the P187A mutation identified in this study did not impact VEEV NHC susceptibility *in vitro* or *in vivo*.

The spatial mapping of the 4’-FlU-associated nsP4 mutations reveals a clustered distribution within the fingers domain, suggesting a shared structural mechanism underlying resistance. Specifically, P187 and Q191 reside in the index finger domain, while L289 and T296 are located in the ring finger domain. These residues form a compact cluster across the GDD active site within the nsP4 core, with inter-residue distances of 7-19 Å, suggesting a coordinated role in nucleoside selection or RNA positioning. While P187 and Q191 have been identified in prior studies, this is the first report linking L289 and T296 to escape from nucleoside analog antivirals. Both substitutions are positioned within a domain previously associated with favipiravir resistance (22) and have been implicated in interactions with incoming nucleotides (36), supporting a potential role in substrate recognition (23).

Viruses can develop resistance to nucleoside analogs through a variety of mechanisms (37–40). The appearance of virus populations with reduced susceptibility to antivirals can be detrimental in the successful implementation of antiviral therapies and the alleviation of virus induced disease. Understanding the susceptibility profiles against antiviral therapeutics is critical for developmental and, ultimately, clinical success. In previous studies, 4’-FlU has been shown to induce immediate or delayed chain termination after incorporation into newly synthesized RNA (25, 26). For some viruses, such as SARS-CoV-2, the ability to proofread has been shown to reduce the incorporation of chain terminators, such as remdesivir (41). While VEEV lacks such a mechanism, alphavirus mutations like CHIKV nsP4-C483Y may enhance polymerase fidelity and confer partial resistance. The absence of this mutation in VEEV under 4’-FlU pressure suggests either limited fitness compatibility, distinct structural constraints, or susceptibility profiling was not saturated.

Despite the relatively rapid emergence of 4’-FlU-associated mutations *in vitro*, 4’-FlU retained significant antiviral activity *in vivo*. Treatment resulted in complete survival and markedly reduced viral titers in mouse models of lethal VEEV encephalitis. Furthermore, viruses encoding multiple nsP4 mutations displayed diminished replication fitness in both cell culture and animal models, suggesting a high genetic barrier preventing robust resistance. These findings support the continued development of 4’-FlU as a promising therapeutic candidate for alphavirus infections.

## Methods Cells

Baby hamster kidney cells (BHK-21; ATCC), African green monkey kidney epithelial cells (VeroE6, CCK-81; ATCC) and VeroE6 cells stably expressing human signaling lymphocytic activation molecule (Vero-hSLAM) or human TMPRSS2 were maintained at 37°C and 5% CO_2_ in Dulbecco’s modified Eagle’s medium (DMEM) supplemented with 8% fetal bovine serum. All immortalized cell lines used in this study are routinely checked for mycoplasma and microbial contamination.

### Molecular Virology

VEEV-TC83 NanoLuciferase (nLuc) stocks were produced from an infectious clone of VEEV-TC83 (generated from VEEV-TC83 and VEEV-TC83-GFP stocks provided by Dr. Ilya Frolov, UAB). Infectious cDNA clones were linearized and purified by gel extraction or using a Qiagen spin column (Qiagen). Capped RNA transcripts were generated using an mMESSAGE mMACHINE SP6 Transcription Kit. *In vitro* transcripts were transfected into BHK-21 cells using Lipofectamine MessengerMax transfection reagent, in accordance with the manufacturer’s protocols. Transfected cells were cultured at 37°C for 24-48 hours until cytopathic effects were observed. Subsequently, supernatants were then harvested and clarified by centrifugation. Virus stocks were aliquoted and titrated by TCID_50_ titration on BHK-21 cells. For recombinant viruses, mutations were generated using site-directed mutagenesis using a helper construct containing the nsP1-4 sequences. After Sanger sequencing confirmed each mutation, mutations were inserted into full-length VEEV-TC83 infectious clones by restriction digest and ligation using T4 DNA ligase. Prior to virus rescue, mutations were confirmed in full-length infectious clones using Sanger sequencing.

### Replicon Assay

An EEE replicon was created from cDNA of EEE nsP1-4 synthesized by Genscript. The EEE replicon cloned into a plasmid and was placed under control of a CMV promoter with NanoLuciferase (nLuc) under control of the EEE subgenomic reporter. BHK-21 cells in 96-well format were transfected with EEE-nLuc replicon plasmid (GeneJuice transfection reagent (Invitrogen)) in accordance with the manufacturer’s protocols (0.1 µg DNA per well). Serial dilutions of compound were added to cells 4 hours after transfection. NanoLuciferase expression was measured 24 hours after transfection.

### Viruses

CHIKV-nLuc, SINV-gLuc, and DENV2-nLuc stocks (provided as a gift from Margaret Kielian and Andres Merits) were propagated on BHK-21 cells and titrated by TCID_50_ titration on BHK-21 cells. Zika-nLuc was provided by Pei-Yong Shi (UTMB) under an MTA. MeV-nLuc stocks were grown on Vero-hSLAM cells (multiplicity of infection (MOI) 0.01 TCID_50_ units/cell). HPIV3-nLuc, VSV-nLuc, and SeV-FLuc (Sendai virus), and SARS-CoV-2-nLuc stocks were grown on VeroE6-TMPRSS2 cells and titered by TCID_50_ titration. Dose-response assays were performed on VeroE6-TMPRSS2 cells, unless otherwise stated.

### Adaptations and susceptibility profiling

VeroE6-TMPRSS2 cells were seeded at 1 × 10^5^ cells/well in 24-well plates, cultured overnight, and inoculated with VEEV-TC83 GFP stock at an MOI of 0.1 TCID_50_/cell. Media containing vehicle (0.1% DMSO) or 4’-FlU (2.5 µM for P1; increasing 2-fold for each passage) was added to each well. After 18-24 hours, supernatant was harvested, divided into aliquots and frozen at -80°C. After each passage, virus was titered by TCID_50_. VeroE6-TMPRSS2 were then infected (MOI=0.1) and incubated with either vehicle or 4’-FlU. Virus lineages were passaged 5-times in the presence of vehicle or 4’-FlU (P5 4’-FlU concentration of 50 µM). After passage 5, RNA was isolated from virus stocks and submitted for whole genome sequencing (Zymo Fast Quick-RNA viral kit (Zymo #R1035).

### Viral next-generation sequencing

Viral whole-genome sequencing of VEEV-TC83 samples was performed using a metagenomic next-generation sequencing approach, as previously described (42). Viral RNA was extracted from cell-culture supernatants using a Zymo Fast Quick-RNA viral kit (Zymo #R1035). Metagenomic sequencing libraries were prepared by Turbo DNAse treatment of RNA extracts (Thermo #AM1907), followed by double-stranded cDNA synthesis using SuperScript IV (Thermo #18090010), random hexamers (Thermo #N8080127), and Sequenase 2.0 (Thermo #70775Z1000UN) and clean-up using 1.8× AMPure XP reagent (Beckman Coulter #A63882), following manufacturers’ recommended protocols. Sequencing libraries were generated from purified cDNA libraries using bead-linked tagmentation using the Illumina DNA Prep with Enrichment Kit (Illumina #20025524) and 14 cycles of PCR amplification using IDT for Illumina DNA/RNA UD Indexes Set D (Illumina #20042667) followed by 1.8 × AMPure XP magnetic bead clean-up. Libraries were sequenced on either NextSeq 2000 or NovaSeq 6000 with 2×150bp read format.

### Data analysis

Raw reads were trimmed and quality filtered with fastp (v0.23.4) (43) with these parameters: --cut_mean_quality 20 --cut_front --cut_tail --length_required 20 --low_complexity_filter --trim poly_g --trim_poly_x. The consensus sequence of VEEV-TC83 GFP P0 stock was generated by mapping the reads to the VEEV TC83 isolate seqeuence (GenBank L01443.1) using Geneious Prime 2024.0. Filtered reads were then used for variant calling using the paired-end RAVA workflow with the VEEV-TC83 GFP P0 stock consensus (GenBank PX275948) as the reference (https://github.com/greninger-lab/RAVA_Pipeline/tree/2025-08-14_GSU_JW_RC_4Flu_VEEV_publication) (44), (https://doi.org/10.1101/2019.12.17.879320).

### Growth Curves

24 hours prior to infection, 1.1 × 10^5^ BHK cells were seeded into each well of a 24-well. Cells were infected with individual VEEV-TC83 strains (MOI = 0.1) which were diluted in serum free media (DMEM). Virus was allowed to adsorb for 1 hour. Subsequently, inoculum was removed, and cells were washed 3 times with complete media (DMEM + 7.5% FBS). Supernatant was harvested at designated time points following infection. Supernatant virus was titrated by TCID_50_. Growth curves were plotted in Prism (Version 10.3).

### Dose response assays

4’-FlU was added in 3-fold dilutions (0.046-100 μM) to 96 well plates seeded with BHK-21 or VeroE6-TMPRSS2 cells (1.5×10^4^/well), followed by infection with HPIV3-JS-nLuc, MeV-nLuc, SeV-Fushimi-GLuc-P2A-eGFP, CedV-nLuc, VSV-nLuc, VEEV-nLuc, SARS-CoV-2-nLuc, CHIKV-nLuc, SINV-Gluc, ZIKV-nLuc (BHK-21), DENV2-nLuc (BHK-21) reporter viruses (MOI = 0.2 TCID_50_ units/cell) and automated plate reading 18-30 hours after infection. Cytotoxic concentrations were determined in equivalent serial dilution plates after exposure of uninfected cells for 72 hours to 4’-FlU, followed by addition of PrestoBlue substrate (Invitrogen) to quantify cell metabolic activity as described (45). Cycloheximide and DMSO vehicle (0.1% DMSO) were used as positive and negative controls, respectively. Four-parameter variable slope regression was used to determine EC_50_, EC_90_, and CC_50_ concentrations.

### Animal studies

For efficacy studies, C3H/HeN mice were infected with the engineered, recombinant VEEV-TC83 variants or the genetic parent virus (VEEV-TC83), administered intranasally at 1.0×10^6^ TCID_50_ units in 50 μl of PBS. Treatment with 4’-FlU (5 mg/kg; q.d.) or molnupiravir (150 mg/kg; b.i.d) commenced 12 hours after infection and continued for seven days. 4’-FlU was administered per oral gavage in a once daily (q.d.) regimen. Molnupiravir was administered in a twice daily regimen (b.i.d) Infected mice were monitored twice daily for clinical signs (body weight loss, overall composure) and euthanized upon reaching endpoint or loosing >20% of their initial weight. Brain viral titers were determined in groups of mice on day 4 after infection through TCID_50_ titration of tissue homogenates on BHK-21 cells.

### Ethics statement

All animal studies were performed following the *Guide for the Care and Use of Laboratory Animals*. All experiments were approved by the Institutional Animal Care and Use Committee of Georgia State University (protocol A24042) and conducted in compliance with the Association for the Accreditation of Laboratory Animal Care guidelines, National Institutes of Health regulations, Georgia State University policy, and local, state, and federal laws.

## Data availability

Raw sequencing data are publicly available under NCBI BioProject accession number PRJNA1306184. Data standing behind figures may be made available upon request to RMC.

## Acknowledgments

We thank S. Atasheva for critical reading. This work was supported, in part, by a Georgia State University Research Initiation Grant (to RMC). The funders had no role in study design, data collection and interpretation, or the decision to submit the work for publication.

